# The ghrelin receptor GHSR has two efficient agonists in an ancient fish species

**DOI:** 10.1101/2023.06.03.543543

**Authors:** Hao-Zheng Li, Ya-Fen Wang, Yong-Shan Zheng, Ya-Li Liu, Zeng-Guang Xu, Zhan-Yun Guo

## Abstract

The gastric peptide ghrelin and its receptor GHSR have important functions in energy metabolism. Recently, liver-expressed antimicrobial peptide 2 (LEAP2) was identified as an endogenous antagonist for GHSR. Ghrelin, LEAP2, and GHSR are ubiquitously present from fishes to mammals and are highly conserved in evolution. However, our recent study suggested that GHSRs from the Actinopterygii fish *Danio rerio* (zebrafish) and *Larimichthys crocea* (large yellow croaker) have lost their binding to ghrelin, despite binding normally to LEAP2. Do these fish GHSRs use another peptide as their agonist? To answer this question, in the present study, we tested to two fish motilins that are closely related to ghrelin. In ligand binding and activation assays, the fish GHSRs from *D. rerio* and *L. crocea* displayed no detectable or very low binding to all tested motilins; however, the GHSR from the Sarcopterygii fish *Latimeria chalumnae* (coelacanth) bound to its motilin with high affinity and was efficiently activated by it. Therefore, it seemed that motilin is not a ligand for GHSR in *D. rerio* and *L. crocea*, but is an efficient agonist for GHSR in *L. chalumnae*, which is known as a ‘living fossil’ and is believed to be one of the closest fish ancestors of tetrapods. The results of present study suggested that in ancient fishes, GHSR had two efficient agonists, ghrelin and motilin; however, this feature might be only preserved in some extant fishes with ancient evolutionary origins. Our present work shed new light on the ligand usage of GHSR in different fish species and in evolution.

## 1. Introduction

The gastric peptide ghrelin is an endogenous agonist for growth hormone secretagogue receptor (GHSR or GHS-R), which can mediate the functions of certain synthetic growth hormone secretagogues (GHSs), hence its name [1,2]. Thus, GHSR is also known as the ghrelin receptor. In some species, the *GHSR* gene can produce two transcripts: one encodes a functional longer isoform 1a (GHSR1a) with a typical architecture of G protein-coupled receptors (GPCRs), while the other encodes a nonfunctional shorter isoform 1b (GHSR1b) lacking a transmembrane domain (TMD). In the present study and in published papers, GHSR refers to the functional longer isoform unless otherwise stated. The mature ghrelin is derived from precursors encoded by the *GHRL* gene, and carries a fatty acyl modification at its third serine residue [1]. This special posttranslational modification is catalyzed by ghrelin *O*-acyltransferase (GOAT), also known as membrane bound *O*-acyltransferase domain containing 4 (MBOAT4) [3,4]. In recent years, liver-expressed antimicrobial peptide 2 (LEAP2 or LEAP-2) has been identified as an endogenous antagonist for GHSR, with low inverse agonist activity [5□7]. LEAP2 was first identified in 2003 and was named according to its weak antimicrobial activity [8]; however its real biological function remained unknown for a long time. The ghrelin-LEAP2-GHSR system has important functions in energy metabolism and cellular homeostasis [9□13].

The genes encoding ghrelin, LEAP2, and GHSR are ubiquitously present from fishes to mammals, implying that they have important functions in vertebrates. Our recent study demonstrated that the antagonistic function of LEAP2 and the agonistic function of ghrelin are conserved in the Sarcopterygii fish *Latimeria chalumnae*, an extant coelacanth believed to be one of the closest fish ancestors of tetrapods [14]. However, GHSRs from the Actinopterygii fish *Danio rerio* (zebrafish) and *Larimichthys crocea* (large yellow croaker) have lost their binding to all tested ghrelins, despite binding normally to LEAP2s [15]. Do these unusual fish GHSRs use another peptide as their agonist? To answer this question, in the present study we tested two fish motilins from *D. rerio* and *L. chalumnae*. Motilin is a gut peptide that stimulates gastrointestinal motility by activating its receptor, MLNR, an A-class G protein-coupled receptor [16□19]. Motilin is closely related to ghrelin, and MLNR is closely related to GHSR; however, human motilin and human ghrelin have no cross activity to each other’s receptor. Herein, we demonstrated that the GHSRs from *D. rerio* and *L. crocea* had no detectable or very low binding to the tested human or fish motilins; however, GHSR from *L. chalumnae* was efficiently activated by its motilin. Thus, it seemed that GHSR in the ancient Sarcopterygii fish *L. chalumnae* has two efficient agonists, ghrelin and motilin, whereas GHSRs from the Actinopterygii fish *D. rerio* and *L. crocea* have lost their binding to both agonists. Our findings provided new insights into the ligand usage of GHSR in different fish species and in evolution.

## 2. Materials and methods

### 2.1. Peptide synthesis

The mature motilins from human or fish were chemically synthesized at GL Biotech (Shanghai, China) via solid-phase peptide synthesis using standard Fmoc methodology. The crude synthetic peptides were purified to homogeneity by high performance liquid chromatography (HPLC) using a semi-preparative C_18_ reverse-phase column (Zorbax 300SB-C18, 9.4 × 250 mm; Agilent Technologies, Santa Clara, CA, USA). Mature human or fish ghrelins and LEAP2s were prepared in our previous studies via chemical synthesis or recombinant expression [14,15].

### 2.2. Preparation of the SmBiT-based tracers

The SmBiT-conjugated LEAP2 tracer, LEAP2-SmBiT, was generated in our previous studies via chemical conjugation of a C-terminally Cys-tagged synthetic SmBiT with a C-terminally Cys-tagged human LEAP2 mutant [21]. The SmBiT-based motilin tracer was prepared according to the procedure for preparation of LEAP2-SmBiT via chemical conjugation of the C-terminally Cys-tagged synthetic SmBiT with a C-terminally Cys-tagged motilin from *L. chalumnae*.

### 2.3. The NanoBiT-based homogenous ligand□receptor binding assays

The expression constructs for the N-terminally secretory large NanoLuc fragment (sLgBiT)-fused human or fish GHSRs were generated in our previous studies [14,15]. These constructs were transiently transfected into human embryonic kidney (HEK) 293T cells, respectively. The next day of transfection, the cells were trypsinized, seeded into white opaque 96-well plates, and continuously cultured for ∼24 h to ∼90% confluence. To conduct the binding assays, the medium was removed and binding solution [phosphate-buffered saline (PBS) plus 0.1% bovine serum albumin (BSA) and 0.01% Tween-20] was added (50 µl/well). For saturation binding assay, the binding solution contained varied concentrations of the SmBiT-based tracer. For competition binding assay, the binding solution contained a constant concentration of a SmBiT-based tracer and varied concentrations of competitor. After incubation at 24 °C for ∼1 h, 20-fold diluted NanoLuc substrate (Promega, Madison, WI, USA) was added (diluted by PBS, 10 µl/well) and bioluminescence was immediately measured on a SpectraMax iD3 plate reader (Molecular Devices, Sunnyvale, CA, USA). The measured bioluminescence data were expressed as mean ± standard deviation (SD, *n* = 3) and fitted to one-site binding model using SigmaPlot 10.0 software (SYSTAT software, Chicago, IL, USA).

### 2.4. Receptor activation assays

The expression constructs for the untagged human or fish GHSRs were generated in our previous studies [14,15], and transiently transfected into HEK293T cells together with a cAMP response element (CRE)-controlled NanoLuc reporter vector pNL1.2/CRE. The next day, the transfected cells were trypsinized, seeded into white opaque 96-well plates, and continuously cultured for ∼24 h to ∼90% confluence. To conduct the activation assays, the medium was removed and activation solution (serum-free DMEM medium plus 1% BSA) was added (50 µl/well). The activation solution contained varied concentrations of ghrelin or motilin. After the cells were continuously cultured at 37 °C for ∼4 h, 20-fold diluted NanoLuc substrate was added (diluted by PBS, 10 µl/well) and bioluminescence was immediately measured on a SpectraMax iD3 plate reader (Molecular Devices). The measured bioluminescence data were expressed as mean ± SD (*n* = 3) and fitted to sigmoidal or linear curves using SigmaPlot 10.0 software (SYSTAT software).

## 3. Results and Discussion

### 3.1. Motilins from fish and higher vertebrates

The motilin gene is widely present in vertebrates, from fishes to mammals. They all encode precursors including an N-terminal signal peptide, a mature peptide, and a C-terminal propeptide (Fig. 1 and Fig. S1). In general, their amino acid sequences are not so conserved in evolution; however, their six N-terminal residues display relatively higher conservation than other residues, consistent with the fact that the N-terminal fragment is important for motilin binding to its receptor [16]. For those fishes with ancient evolutionary origins, including the Chondrichthyes fish *Chiloscyllium plagiosum, Rhincodon typus, Stegostoma fasciatum*, and *Scyliorhinus canicular*, and the Sarcopterygii fish *Protopterus annectens* (lungfish) and *Latimeria chalumnae* (coelacanth), their mature motilins have an aromatic Phe reside at the first position (Fig. 1 and Fig. S1). For mature motilins from higher vertebrates, their first residue is also an aromatic Phe or an aromatic Tyr residue. However, for those Actinopterygii fish with late evolutionary origins, such as *D. rerio*, their mature motilins have a His residue at the first position. An aromatic Phe residue occupies the fourth position in all fish motilins, and in most motilins from amphibians to birds; however, this position is typically occupied by a large aliphatic residue in mammalian motilins (Fig. 1 and Fig. S1). For most mature motilins, their fifth position is occupied by an aromatic Phe residue, and their sixth position is occupied by a Ser or Thr residue.

**Fig. 1.**
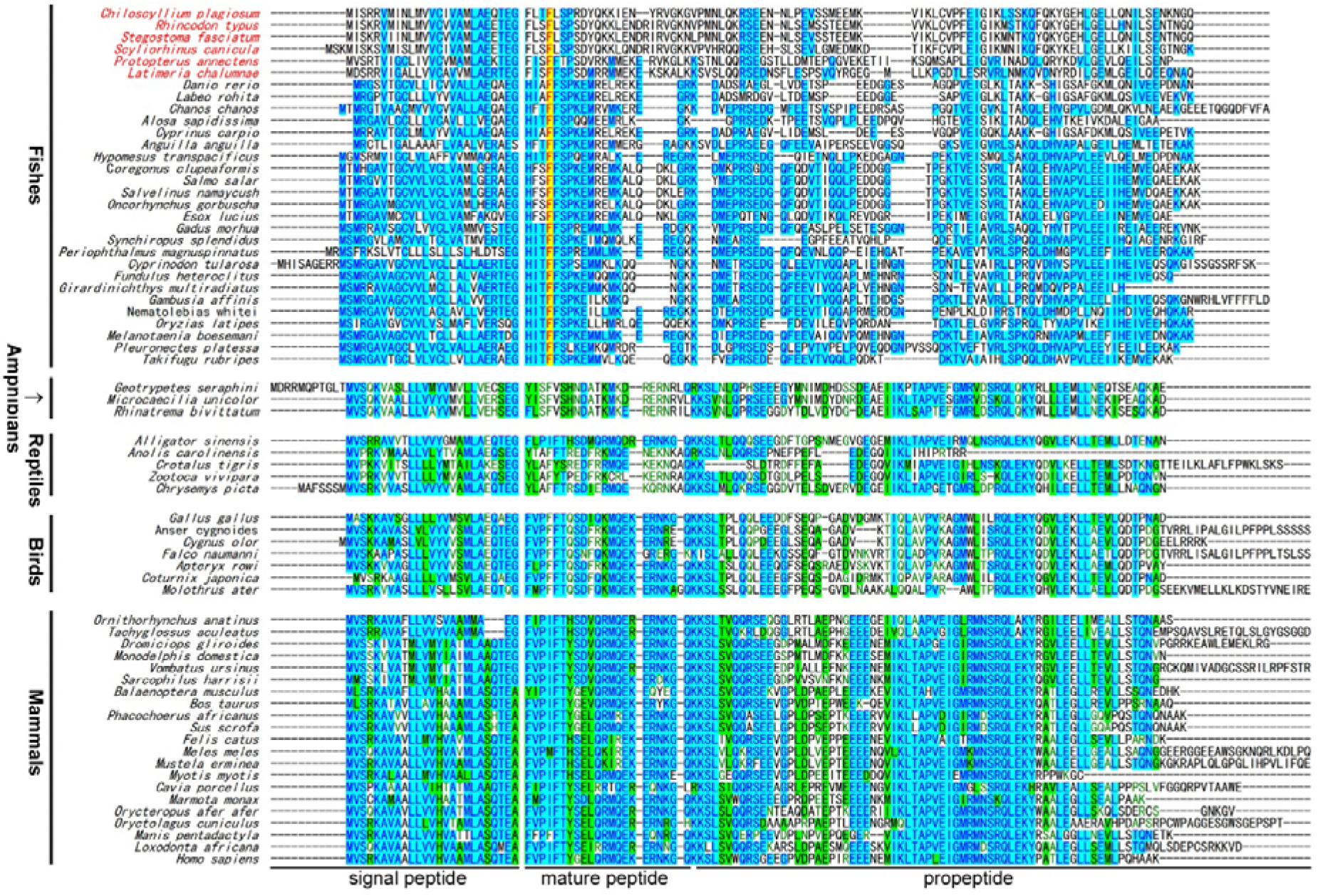
Amino acid sequence alignment of some motilin precursors from fishes to mammals. These sequences were manually downloaded from the NCBI database (https://www.ncbi.nlm.nih.gov/gene) and aligned using the software Vector NTI11.5. The motilin precursor from *L. crocea* was not found in the NCBI database.

To test whether motilin is a ligand of fish GHSRs, we chemically synthesized mature human motilin (Hs-motilin) and two fish motilins, Lc-motilin from *L. chalumnae* and Dr-motilin from *D. rerio* (Fig. 2A). The human motilin gene produces three transcripts (NM_002418, NM_001040109, and NM_001184698) that encode three precursors with slight differences at their C-termini (Fig. S2); however, these precursors release an identical mature peptide after *in vivo* processing. The *D. rerio* motilin-like gene produces two transcripts (NM_001386353 and NM_001386354) that encode two identical precursors (Fig. S2). A Gly residue flanks the C-terminus of the expected mature Dr-motilin (Fig. S2), implying that it has an α-amidated C-terminus. The *L. chalumnae* motilin gene also produces two transcripts (XM_005995467 and XM_014488135) that encode two identical precursors (Fig. S2). The motilin gene from *L. crocea* was not found in the database of National Center for Biotechnology Information (NCBI).

**Fig. 2.**
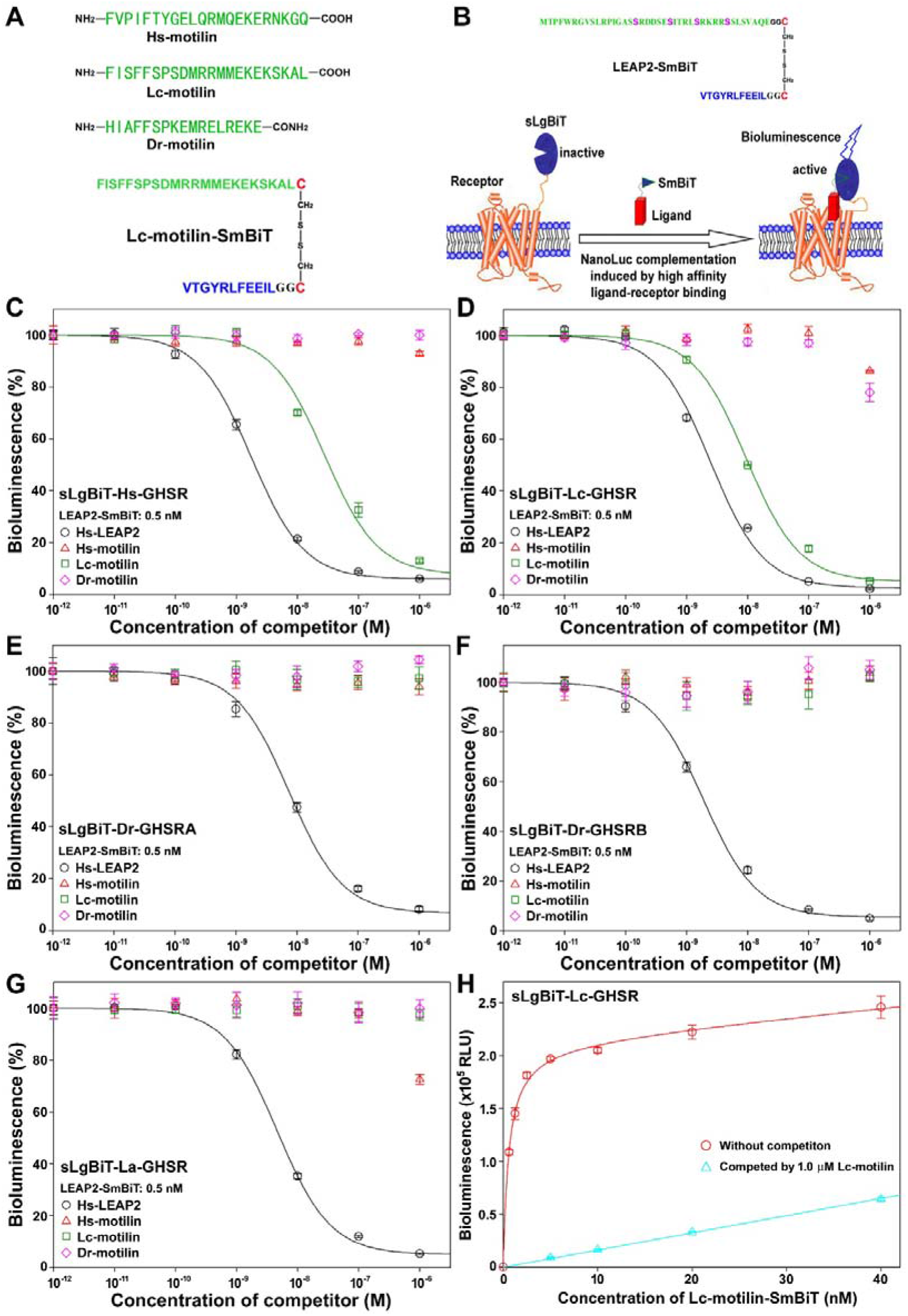
Binding of synthetic motilins to human or fish GHSRs. (**A**) Amino acid sequence of the synthetic motilins and SmBiT-based Lc-motilin tracer. (**B**) Schematic presentation of the NanoBiT-based binding assay. (**C** □ **F**) Competition binding of motilins with the N-terminally sLgBiT-fused Hs-GHSR (**C**), Lc-GHSR (**D**), Dr-GHSRA (**E**), Dr-GHSRB (**F**), and La-GHSR (**G**) using LEAP2-SmBiT as a tracer. (**H**) Saturation binding of Lc-motilin-SmBiT with the N-terminally sLgBiT-fused Lc-GHSR.

### 3.2. Binding of synthetic motilins to human or fish GHSRs

To test whether the synthetic motilins bind to human or fish GHSRs, we employed a NanoBiT-based competition binding assay developed in our previous studies [14,15]. This binding assay relies on an N-terminally sLgBiT-fused receptor and a SmBiT-fused tracer (Fig. 2B). Our previous studies demonstrated that a SmBiT-conjugated human LEAP2 mutant, LEAP2-SmBiT (Fig. 2B), could bind to human or fish GHSRs [14,15], thus we used it as a tracer to measure the receptor-binding potency of the synthetic motilins.

The synthetic Hs-motilin had no detectable binding towards the N-terminally sLgBiT-fused human GHSR (Hs-GHSR) (Fig. 2C), confirming that motilin has no cross-activity with GHSR in human. The synthetic Dr-motilin also had no detectable binding with Hs-GHSR (Fig. 2C). However, the synthetic Lc-motilin could bind to Hs-GHSR, although its binding potency was approximately16-fold lower than that of human LEAP2 (Hs-LEAP2), the endogenous antagonist of Hs-GHSR (Fig. 2C and Table 1).

**Table 1.** Summary of the measured IC_50_ (nM) and EC_50_ (nM) values of motilin, ghrelin, and LEAP2 to human or fish GHSRs.

| Peptides | GHSRs |  |  |  |  |
| --- | --- | --- | --- | --- | --- |
|  | Hs-GHSR | Lc-GHSR | Dr-GHSRA | Dr-GHSRB | La-GHSR |
| Hs-motilin | N.D. | low | N.D. | N.D. | low |
| Lc-motilin | 27.5 ± 2.4<br>[~1.5]<sup>a,b</sup> | 9.55 ± 0.45<br>[2.14 ± 0.26]<sup>a</sup> | N.D. | N.D. | N.D. |
| Dr-motilin | N.D. | low | N.D. | N.D. | N.D. |
| Hs-ghrelin | 3.63 ± 0.24<sup>c</sup><br>[1.74 ± 0.26] | 12.3 ± 1.1<sup>c</sup><br>[4.07 ± 0.52]<sup>c</sup> | N.D.<sup>c</sup> | N.D.<sup>c</sup> | N.D.<sup>c</sup> |
| Lc-ghrelin | 12.0 ± 0.6<sup>c</sup><br>[10.0 ± 1.5]<sup>c</sup> | 6.03 ± 0.41<sup>c</sup><br>[2.63 ± 0.39] | N.D.<sup>c</sup> | N.D.<sup>c</sup> | N.D.<sup>c</sup> |
| Dr-ghrelin-22 | 13.2 ± 0.9<sup>c</sup><br>[6.31 ± 0.94]<sup>c</sup> | 11.7 ± 1.0<sup>c</sup><br>[5.50 ± 0.71]<sup>c</sup> | >1000<sup>c</sup> | ~1000<sup>c</sup> | N.D.<sup>c</sup> |
| La-ghrelin-23 | 6.61 ± 0.31<sup>c</sup><br>[5.37 ± 0.90]<sup>c</sup> | 35.5 ± 1.7<sup>c</sup><br>[6.03 ± 0.41]<sup>c</sup> | N.D.<sup>c</sup> | ~1000<sup>c</sup> | N.D.<sup>c</sup> |
| Hs-LEAP2 | 1.74 ± 0.08 | 2.45 ± 0.16 | 7.41 ± 0.53 | 1.91 ± 0.13 | 4.68 ± 0.21 |
| Lc-LEAP2 | 4.79 ± 0.46<sup>c</sup> | 2.00 ± 0.09<sup>c</sup> | 4.37 ± 0.20<sup>c</sup> | 2.00 ± 0.09<sup>c</sup> | 2.40 ± 0.36<sup>c</sup> |
| Dr-LEAP2 | 23.4 ± 2.5<sup>c</sup> | 9.77 ± 0.65<sup>c</sup> | 5.75 ± 0.28<sup>c</sup> | 3.09 ± 0.14<sup>c</sup> | 3.80 ± 0.25<sup>c</sup> |
| La-LEAP2A | 6.17 ± 0.42<sup>c</sup> | 4.17 ± 0.19<sup>c</sup> | 2.69 ± 0.12<sup>c</sup> | 1.32 ± 0.06<sup>c</sup> | 2.14 ± 0.14<sup>c</sup> |

N.D., not detectable; low, inhibitory effect was only observed at the concentration of 1.0 μM.
<sup>a</sup> EC<sub>50</sub> values are listed in brackets.
<sup>b</sup> E<sub>max</sub> of Lc-motilin activating Hs-GHSR is much lower than that of Hs-ghrelin.
<sup>c</sup> data cited from our previous study [15].

Lc-motilin bound to the sLgBiT-fused GHSR from *L. chalumnae* (Lc-GHSR) with a calculated IC_50_ value of 9.55 ± 0.45 nM (*n* = 3), which was approximately 4-fold higher than that of Hs-LEAP2 (Fig. 2D, Table 1). Our previous studies demonstrated that Hs-LEAP2 displayed approximately 3-fold higher binding potency than Lc-ghrelin towards Lc-GHSR (Table 1) [14,15]. Thus, it seemed that motilin and ghrelin displayed similar binding potency towards GHSR in the Sarcopterygii fish *L. chalumnae*. By contrast, the binding potency of Hs-motilin and Dr-motilin towards Lc-GHSR was too low to be quantified (Fig. 2D and Table 1).

All tested motilins had no detectable binding towards two GHSRs from *D. rerio*, Dr-GHSRA and Dr-GHSRB (Fig. 2E,F). Only Hs-motilin displayed detectable binding towards the fish GHSR from *L. crocea* (La-GHSR); however, it was too low to be quantified (Fig. 2G). Thus, it seemed that motilin displays diverse binding properties with GHSRs in different fish species.

To confirm tight binding of Lc-motilin to Lc-GHSR, we prepared a SmBiT-based tracer for Lc-motilin via chemical conjugation of a C-terminally Cys-tagged SmBiT with a C-terminally Cys-tagged Lc-motilin (Fig. 2A). The resultant Lc-motilin-SmBiT tracer bound to the sLgBiT-fused Lc-GHSR in a saturable manner (Fig. 2H), with a calculated dissociation constant (K_d_) of 0.58 ± 0.04 nM (*n* = 3). Thus, Lc-motilin bound to Lc-GHSR with high affinity, implying that motilin might be an endogenous ligand of GHSR in the ancient fish *L. chalumnae*.

### 3.3. Activation of GHSRs by the coelacanth motilin

Lc-motilin bound to Hs-GHSR and Lc-GHSR; therefore, we tested whether it functions as an agonist or an antagonist via a receptor activation assay using a CRE-controlled NanoLuc reporter. As a positive control, Hs-ghrelin efficiently activated the untagged Hs-GHSR (Fig. 3A and Table 1), with a calculated EC_50_ value of 1.74 ± 0.26 nM (*n* = 3). The synthetic Lc-motilin also activated Hs-GHSR, but its maximal effect (E_max_) was much lower than that of Hs-ghrelin (Fig. 3A and Table 1), suggesting it is not an efficient agonist towards the human receptor. Towards the untagged Lc-GHSR, Lc-ghrelin and Lc-motilin displayed similar EC_50_ values and similar E_max_ values (Fig. 3B and Table 1). Thus, it seemed that motilin is an efficient agonist for GHSR in the ancient Sarcopterygii fish *L. chalumnae*.

**Fig. 3.**
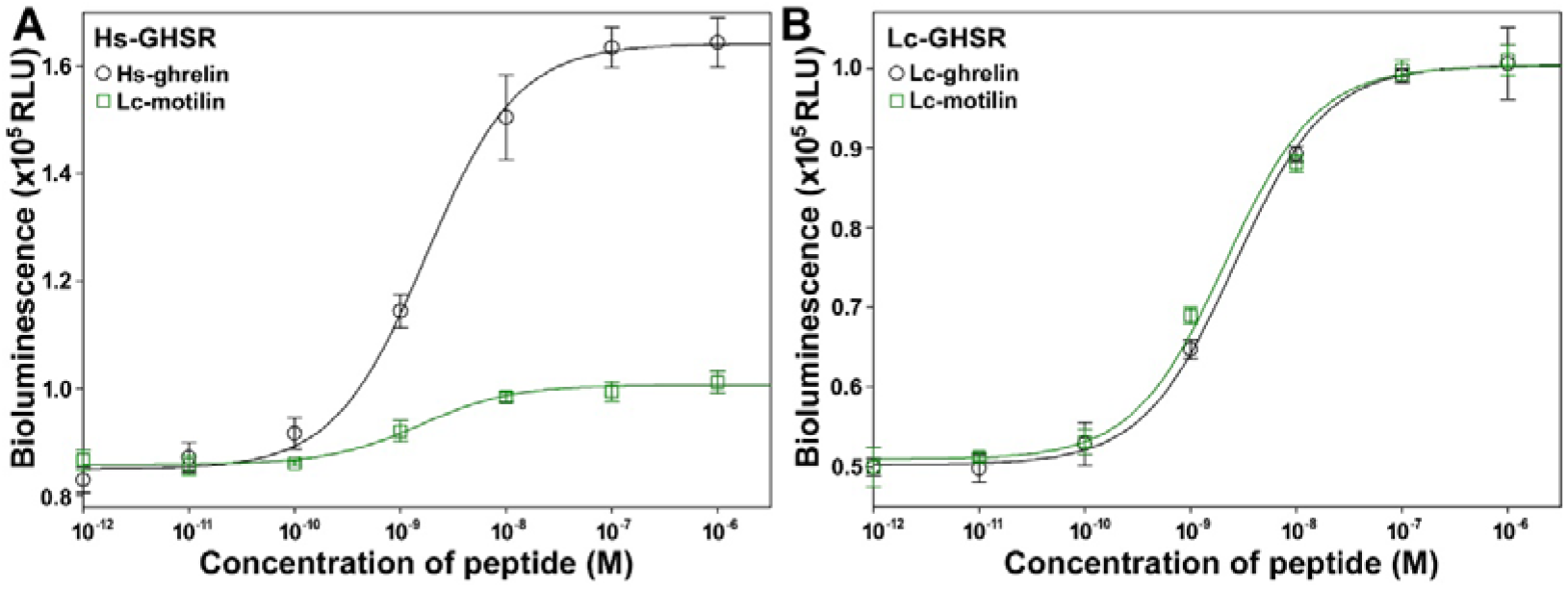
Activation of the untagged Hs-GHSR (**A**) and Lc-GHSR (**B**) by ghrelin and Lc-motilin monitored using a CRE-controlled NanoLuc reporter.

### 3.4. Ligand usage of GHSR in different fish species and in evolution

Our present study suggested that motilin is an efficient agonist for the ghrelin receptor GHSR in the Sarcopterygii fish *L. chalumnae*, an extent coelacanth with an ancient evolutionary origin. Fossil records suggested that coelacanths first appeared in the Devonian era about 400 million years ago, reaching their peak in the Triassic era, and might have been extinct since the Late Cretaceous era about 70 million years ago. However, a living coelacanth specimen was discovered in South Africa in 1938, and this extant coelacanth species was named *Latimeria chalumnae*. To date, only two extant coelacanths (*L. chalumnae* and *L. menadoensis*) have been found in small areas of the Indian Ocean. They are remarkably similar to their ancient relatives, and genomic sequencing disclosed that their genes evolved significantly more slowly than those of tetrapods [22,23]. Thus, it seemed that *L. chalumnae* might retain some ancient features that appeared in the early stage of fish evolution.

The genes encoding ghrelin, LEAP2, GHSR, and motilin are widely present in fishes, but they are absent in lower chordates, such as Branchostoma and Ascidiacea, which suggested they might have originated in ancient fish and then spread to all vertebrate lineages. As shown in Fig. 1 and Fig. S1, motilins from fish with ancient evolutionary origins, including the Chondrichthyes fish *C. plagiosum, R. typus, S. fasciatum*, and *S. canicular*, and the Sarcopterygii fish *P. annectens* and *L. chalumnae*, share similar features, such as an aromatic Phe residue at the first position of their mature peptides. Thus, two efficient GHSR agonists, ghrelin and motilin, might have originated in and evolved from ancient fishes; however, this ancient feature might only be retained in the extant Chondrichthyes and Sarcopterygii fishes with ancient evolutionary origins. Motilins from Actinopterygii fishes are quite different from those from Chondrichthyes and Sarcopterygii fishes, such as a His residue, rather than an aromatic Phe, at the first position of their mature peptides (Fig. 1 and Fig. S1). As a result, it seemed that motilin might be no longer a ligand of GHSR in these ray-finned fishes with much later evolutionary origins. Some features of the ancient fish motilins, such as an aromatic residue at the first position of their mature peptide, were passed down to higher vertebrates (Fig. 1 and Fig. S1). However, motilin is no longer a ligand of GHSR in humans and probably in other mammals [24]. Currently, it is largely unknown whether motilin is a ligand of GHSR in birds, reptiles, and amphibians. Thus, the relationship between motilin and GHSR in different vertebrate species needs to be further tested in future studies.

Our binding and activation assays were conducted on transfected HEK293T cells, thus we cannot exclude the possibility that the overexpressed Actinopterygii fish GHSRs had some defects, such as lack of special posttranslational modifications or lack of interactions with other receptors or proteins, and thus lost binding to their ghrelin and motilin. However, this possibility was not high, considering that the overexpressed receptors retain normal binding to LEAP2. Moreover, the overexpressed fish GHSR from *L. chalumnae* displayed normal binding to its ghrelin, LEAP2, and motilin. In the future, more studies are needed to clarify whether these Actinopterygii fish GHSRs can be activated by ghrelin and motilin *in vivo*.

## Acknowledgements

This work was supported by grant from the National Natural Science Foundation of China (grant no. 31971193).

## Notes

### Competing Interest Statement

The authors have declared no competing interest.

## References

1. Kojima M, Hosoda H, Date Y, Nakazato M, Matsuo H, Kangawa K (1999) Ghrelin is a growth-hormone-releasing acylated peptide from stomach. Nature 402: 656–660.

2. Howard AD, Feighner SD, Cully DF, Arena JP, Liberator PA, Rosenblum CI, Hamelin M, Hreniuk DL, Palyha OC, Anderson J, Paress PS, Diaz C, Chou M, Liu KK, McKee KK, Pong SS, Chaung LY, Elbrecht A, Dashkevicz M, Heavens R, Rigby M, Sirinathsinghji DJ, Dean DC, Melillo DG, Patchett AA, Nargund R, Griffin PR, DeMartino JA, Gupta SK, Schaeffer JM, Smith RG, Van der Ploeg LH (1996) A receptor in pituitary and hypothalamus that functions in growth hormone release. Science 273: 974–977

3. Yang J, Brown MS, Liang G, Grishin NV, Goldstein JL. (2008) Identification of the acyltransferase that octanoylates ghrelin, an appetite-stimulating peptide hormone. Cell 132: 387–396.

4. Gutierrez JA, Solenberg PJ, Perkins DR, Willency JA, Knierman MD, Jin Z, Witcher DR, Luo S, Onyia JE, Hale JE. (2008) Ghrelin octanoylation mediated by an orphan lipid transferase. Proc. Natl. Acad. Sci. U.S.A. 105: 6320–6325.

5. Ge X, Yang H, Bednarek MA, Galon-Tilleman H, Chen P, Chen M, Lichtman JS, Wang Y, Dalmas O, Yin Y, Tian H, Jermutus L, Grimsby J, Rondinone CM, Konkar A, Kaplan DD. (2018) LEAP2 Is an Endogenous Antagonist of the Ghrelin Receptor. Cell Metab 27: 461–469.

6. Wang JH, Li HZ, Shao XX, Nie WH, Liu YL, Xu ZG, Guo ZY. (2019) Identifying the binding mechanism of LEAP2 to receptor GHSR1a. FEBS J 286: 1332–1345.

7. M’Kadmi C, Cabral A, Barrile F, Giribaldi J, Cantel S, Damian M, Mary S, Denoyelle S, Dutertre S, Péraldi-Roux S, Neasta J, Oiry C, Banères JL, Marie J, Perello M, Fehrentz JA. (2019) N-Terminal Liver-Expressed Antimicrobial Peptide 2 (LEAP2) Region Exhibits Inverse Agonist Activity toward the Ghrelin Receptor. J Med Chem 62: 965–973.

8. Krause A, Sillard R, Kleemeier B, Klüver E, Maronde E, Conejo-García JR, Forssmann WG, Schulz-Knappe P, Nehls MC, Wattler F, Wattler S, Adermann K. (2003) Isolation and biochemical characterization of LEAP-2, a novel blood peptide expressed in the liver. Protein Sci 12: 143–152.

9. Abizaid A, Hougland JL. (2020) Ghrelin Signaling: GOAT and GHS-R1a Take a LEAP in Complexity. Trends Endocrinol Metab 31: 107–117.

10. Al-Massadi O, Müller T, Tschöp M, Diéguez C, Nogueiras R. (2018) Ghrelin and LEAP-2: Rivals in Energy Metabolism. Trends Pharmacol Sci 39: 685–694.

11. Cervone DT, Lovell AJ, Dyck DJ. (2020) Regulation of adipose tissue and skeletal muscle substrate metabolism by the stomach-derived hormone, ghrelin. Curr Opin Pharmacol 52: 25–32.

12. Yanagi S, Sato T, Kangawa K, Nakazato M. (2018) The Homeostatic Force of Ghrelin. Cell Metab 27: 786–804.

13. Reich N, Hölscher C. (2022) Beyond appetite: Acylated ghrelin as a learning, memory and fear behavior-modulating hormone. Neurosci Biobehav Rev 143: 104952.

14. Li HZ, Shou LL, Shao XX, Li N, Liu YL, Xu ZG, Guo ZY. (2021) LEAP2 has antagonized the ghrelin receptor GHSR1a since its emergence in ancient fish. Amino Acids 53: 939–949.

15. Li HZ, Shao XX, Wang YF, Liu YL, Xu ZG, Guo ZY. (2023) LEAP2 is a more conserved ligand than ghrelin for fish GHSRs. Biochimie 209: 10–19.

16. Feighner SD, Tan CP, McKee KK, Palyha OC, Hreniuk DL, Pong SS, Austin CP, Figueroa D, MacNeil D, Cascieri MA, Nargund R, Bakshi R, Abramovitz M, Stocco R, Kargman S, O’Neill G, Van Der Ploeg LH, Evans J, Patchett AA, Smith RG, Howard AD. (1999) Receptor for motilin identified in the human gastrointestinal system. Science 284: 2184–2188.

17. Mori H, Verbeure W, Tanemoto R, Sosoranga ER, Jan Tack. (2023) Physiological functions and potential clinical applications of motilin. Peptides 160: 170905.

18. Kitazawa T, Kaiya H. (2021) Motilin Comparative Study: Structure, Distribution, Receptors, and Gastrointestinal Motility. Front Endocrinol (Lausanne) 12: 700884.

19. Singaram K, Gold-Smith FD, Petrov MS. (2020) Motilin: a panoply of communications between the gut, brain, and pancreas. Expert Rev Gastroenterol Hepatol 14: 103–111.

20. Deloose E, Verbeure W, Depoortere I, Tack J. (2019) Motilin: from gastric motility stimulation to hunger signalling. Nat Rev Endocrinol 15: 238–250.

21. Li HZ, Shou LL, Shao XX, Liu YL, Xu ZG, Guo ZY. (2020) Identifying key residues and key interactions for the binding of LEAP2 to receptor GHSR1a. Biochem J 477: 3199–3217.

22. Amemiya CT, Alföldi J, Lee AP, Fan S, Philippe H, Maccallum I, Braasch I, Manousaki T, Schneider I, Rohner N, Organ C, Chalopin D, Smith JJ, Robinson M, Dorrington RA, Gerdol M, Aken B, Biscotti MA, Barucca M, Baurain D, Berlin AM, Blatch GL, Buonocore F, Burmester T, Campbell MS, Canapa A, Cannon JP, Christoffels A, De Moro G, Edkins AL, Fan L, Fausto AM, Feiner N, Forconi M, Gamieldien J, Gnerre S, Gnirke A, Goldstone JV, Haerty W, Hahn ME, Hesse U, Hoffmann S, Johnson J, Karchner SI, Kuraku S, Lara M, Levin JZ, Litman GW, Mauceli E, Miyake T, Mueller MG, Nelson DR, Nitsche A, Olmo E, Ota T, Pallavicini A, Panji S, Picone B, Ponting CP, Prohaska SJ, Przybylski D, Saha NR, Ravi V, Ribeiro FJ, Sauka-Spengler T, Scapigliati G, Searle SM, Sharpe T, Simakov O, Stadler PF, Stegeman JJ, Sumiyama K, Tabbaa D, Tafer H, Turner-Maier J, van Heusden P, White S, Williams L, Yandell M, Brinkmann H, Volff JN, Tabin CJ, Shubin N, Schartl M, Jaffe DB, Postlethwait JH, Venkatesh B, Di Palma F, Lander ES, Meyer A, Lindblad-Toh K. (2013) The African coelacanth genome provides insights into tetrapod evolution. Nature 496: 311–316

23. Nikaido M, Noguchi H, Nishihara H, Toyoda A, Suzuki Y, Kajitani R, Suzuki H, Okuno M, Aibara M, Ngatunga BP, Mzighani SI, Kalombo HW, Masengi KW, Tuda J, Nogami S, Maeda R, Iwata M, Abe Y, Fujimura K, Okabe M, Amano T, Maeno A, Shiroishi T, Itoh T, Sugano S, Kohara Y, Fujiyama A, Okada N. (2013) Coelacanth genomes reveal signatures for evolutionary transition from water to land. Genome Res 23: 1740–1748.

24. Nunoi H, Matsuura B, Utsunomiya S, Ueda T, Miyake T, Furukawa S, Kumagi T, Ikeda Y, Abe M, Hiasa Y, Onji M. (2012) A relationship between motilin and growth hormone secretagogue receptors. Regul Pept 176: 28–35.

