## Supplementary material for "The ghrelin receptor GHSR has two efficient agonists in an ancient fish species": Fig. S1-S2

**Contents:**

**Fig. S1.** Amino acid sequence alignment of motilin precursors from fishes to mammals.

**Fig. S2.** Amino acid sequence of motilin precursors encoded by transcript variants from *H. sapiens*, *D. rerio*, and *L. chalumnae.*

```
Alosa alosa                   (1) ------------MRGAVLGCLLLVCAVLLLVEQAEG HITFFSPQQMEEMRLK------GK-----GPRSEDK-TPEETSVQPLPLEEDPQV----HGTEVEISIKLTADQLEHVTKEIVKDALEIGAA---------------
Alosa sapidissima             (1) ------------MRGAVLGCLLLVCAVLLLVEQAEG HITFFSPQQMEEMRLK------GK-----GPRSEDK-TPEETSVQPLPLEEDPQV----HGTEVEISIKLTADQLEHVTKEIVKDALEIGAA---------------
Carassius gibelio             (1) ------------MRRAVTRCLMLVCVVALLAEQAEG HIAFFSPKEMRELKEKE-----GRR--DADLRAEGV-FIDEMSP----DGD--ES----VGQPVEIGLKLTAKK-GQIGSAFGKMLQSIIEEPETVK-----------
Cyprinus carpio               (1) ------------MRRAVTGCLMLVYVVALLAEQAEG HIAFFSPKEMRELREKE-----GRK--DADPRAEGV-LIDEMSL----DEE--ES----VGQPVEIGQKLAAKK-GHIGSAFDKMLQSIVEEPETVK-----------
Puntigrus tetrazona           (1) ------------MRRAVTGCLMLVCVVALLADQAEG HIAFFSPKEMRELREKE-----GRK--DADPRAEGV-LIEETSL----DEDGGES----AGQPVEIGLKLTAKK-GHIGSAFGKMLQSIVEEPETAK-----------
Megalobrama amblycephala      (1) ---- -------MRGAVTGCLVLVYVIALQADQAEG HIAFFSPKEMRELREKE-----GRK--DADLRAEEV-LIDEMSP----EENRGES----AGQLVEIGLKLTAKK-GHIGSAFGKMLQNIVEEPETVK-----------
Pimephales promelas           (1) ------------MRVAVTGCLVLLYIVALQGDQADG HITFFSPKEMRELREKE-----GRK--DADPQAVEV-LIDETG---------GEP----VGQPIEIGVKLTAKK-GHIGSAFGKMLQNIVGEPEMVK-----------
Danio rerio                   (1) ------------MRGSVTGCVLLICVVALLAEQAEG HIAFFSPKEMRELREKE-----GRK--DADSRAEGL-LVDETSP----EEDGGES----AGQPVEIGLKLTAKK-SHIGSAFGKMLQNIVEEPDNAN-----------
Labeo rohita                  (1) ------------MRGPVTGCLLLVYVVALLAEQAEG HIAFFSPKEMRELREKE-----GRK--DADSMRDGV-LTDEMSP----EEDGGE------SAPVEIGLKLTAKK-GHIGSAFGKMLQSIVEEVEKVK-----------
Chanos chanos                 (1) ----------MTMRGTVAACMVVCVALLAEQAEG HFTFFSPKEMRVMKERE-----GKK--DVEPRSEDG-MFEETSVSPIPEEDRSAS----PGQTVEIGVKLTAKQLEHVGPVLGDMLQKVLNEAEKGEEETQGQDFVFA
Chiloscyllium plagiosum       (1) -----------MISRRVMINLMVVCIVAMLAEQTEG FLTFLSPRDYQKKIEN--YRVGKGVPMNLQKRSEEN-NLPEVSSMEEMK-------VIKLCVPFEIGIKLSSKQFQKYGEHLGELLQNILSENKNGQ-----------
Rhincodon typus               (1) -----------MISKRVMINLMVVCVVAMLAEETEG FLSFLSPSDYQKKLENDRIRVGKNVPMNLQKRSEEN-NLSEMSSTEEMK-------VVKLCVPFEIGIKMSTKQFQKYGEHLGELLHNILSENTNGQ-----------
Stegostoma fasciatum          (1) -----------MISKRVIINLMVVCVVAMLAEETEG FLSFLSPSDYQKKLENDRIRVGKNLPMNLQKRSEEN-NLSEVSSTEEMK-------VIKLCVPFEIGIKMNTKQYQKYGEHLGELLQNILSENTNGQ-----------
Scyliorhinus canicula         (1) --------MSKMISKSVMISLMVVCIVAMLAEETEG FLSFLSPSDYQKKLQNDRIRVGKKVPVHRQQRSEEH-SLSEVLGMEDMKD------TIKFCVPLEIGIKMNIKQFQKYKELLGELLKIILSEGTNGK-----------
Protopterus annectens         (1) -----------MVSRTVIGCLIVVCVMAMLAEKTEG FISFFTPSDVRKMMEKE-RVKGLKKSTNLQQRSEGSTLLDMTEPQGVEKETII---KSQMSAPLEIGVRINADQLQRYKDVLGEVLQEILSENP-----------
Latimeria chalumnae           (1) -----------MDSRRVIGALLVVCAVVMLAERTEG FISFFSPSDMRRMMEKE-KSKALKKSVSLQQRSEDNSFLESPSVQYRGEG--M---LLKPGDTLESRVRLNMKQVDNYRDILGEMLGEILQEEQNAQ-----------
Anguilla anguilla             (1) ------------MRCTLIGALAAAFLVAALVERAES HFTFFSPKEMREMMERG---RAGKKSVDLEPRSEEG-QFEEVAIPERSEEVGGSQ-----GKSVEIGVRLSAKQLDHVAPALGEILHEMLTETEKAK-----------
Hypomesus transpacificus      (1) ----------MGMSRMVIGCLVLAFFVVMMAQRAEG HITFFSPQEMRALK--E---REGRK--LMEPRSEDG---QIETNQLLPKEDGAGN----PEKTVEISMQLSAKQLDHVAPVLEEVLQELMEDPDNAK-----------
Coregonus clupeaformis        (1) ----------MTMHGAVTGCVVLVCLVAMLGERAEG HFSFFSPKEMREMKALQ--DKLGRK--DMKPRSGDG-QFQDVTIQQLPEDDGG-----TPEKTVEISVRLTAKQLEHVAPVLEEIIHEMVEQAEKKAK-----------
Salmo salar                   (1) ----------MTMRGVVTGCVVLVCLVAMLGERAEG HFSFFSPKEMREMKALQ--DKLGRK--YMEPRSEDG-QFQDVTIQQLPEDDGG-----TPGKTVEISVRLTAKQLEHVAPVLEEIIHEMVEQAEEKAK-----------
Salvelinus namaycush          (1) ----------MTMRGAVTGCVVLVCLVAMLGERAEG HFSFFSPKEMREMKALQ--DKLERK--DMEPRSEDG-QFQDVTIQQLPEDDGG-----TPGKTVEISVRLTAKQLEHVAPVLEEIIHEMVDQAEKKAK-----------
Oncorhynchus gorbuscha        (1) ----------MTMRGAVMGCVVLVCLVAMLHERAEG HFSFFSPKEMRELKALQ--DKLGRK--DMEPRSEDG-QFQDVTIQQLPEDDGG-----TPGKTVEISVRLTAKQLEHVAPVLEDIIHEMVEQAEKKAK-----------
Oncorhynchus keta             (1) ----------MTMRGAVTGCVVLVCLVAMLHERAEG HFSFFSPKEMRELKALQ--DKLGRK--DMEPRSEDG-QFQDVTIQQLPEDDGG-----TPGKTVEISVRLTAKQLEHVAPVLEDIIHEMVEQAEKKAK-----------
Oncorhynchus mykiss           (1) ----------MTMRGAMTGCVVLVCLVAMLHERAEG HFSFFSPKEMREMKALQ--NKLGRK--DMEPRSEDG-QFQDVTIQQLPEDDGG-----TPGKTVEISVRLTAKQLEHVAPVLEDIIHEMVEQAEKKAK-----------
Esox lucius                   (1) ----------MTMRGAVMCCVLLVCLVAMFAKQVEG HFSFFSPKEMREMKALQ--NKLGRK--DMEPQTENG-QLQDVTIKQLREVDGR-----IPEKIMEIGVRLTAKQLELVGPVLEEIINEMVEQAE-----------
Gadus morhua                  (1) ----------MSMRRAVSGCVLVVCLVAMMVESTEG HITFFSPREMMLMK--E---RDGKK--VMEPRSEDG-QFQEASLPELSETESGGN----PDRTIEIAVRLSAQQLYHVTPVLEEIIREIAEEREKVNK-----------
Synchiropus splendidus        (1) ----------MSMRGVLAMCVVLTCLVATMVERTEG HITFFSPKEIMQMQLKE---REGQK--NMEARSE------EGPFEEATVQHLP------QDETVEIAVRLSPQQLDHVAPVLEEIIHQIAGENRKGIRF----------
Periophthalmus magnuspinnatus (1) --------MRMSFRKSLVTCLLLSLLLSLHLDTSEG HITFFSPREMMLMK--E---REGKK--DMEPRSEDG-QFQEVNLQQP-EIEHGAT----PEKAVEVTVRLSPRQLDHMGPVLEEFIHEIVEERQKAK-----------
Cyprinodon tularosa           (1) -MHISAGERRMSMRGAVVGCVVLVCLLALLAERTEG HITFFSPSEMMKLKQQ-----NGKK--NMETRSEDG-QLEEVTVQQAPLIEHNGN----PDNTLEVAIRLLPRQVDHVSPVLEEIIHEIVEQSQKGISSGSSRFSK--
Fundulus heteroclitus         (1) ----------MSMRGAVAGCVVLACLLALVAERTEG HITFFSPKEMQQMKQQ-----NGKK--DMETRSEDG-QFEEVTVQQAPLMEHNRN----SDNTLEVAVRLLPRQVDHVAPVLEEIIHEIVEQSQ-------------
Girardinichthys multiradiatus (1) ----------MSMRRAVAGCVVLMCLLALVAERTEG HITFFSPKEMMKMKQQ-----NGKK--DMETRSEDG-QFEEVIVQQAPLVEHNGN----SDN-TEVAVLLLPRQMDQVPPALEEILH----------------------
Gambusia affinis              (1) ----------MSMRGAVAGCVVLVCLLALVVERTEG HITFFSPKEILKMKQ------NGKK--DMEARSEDG-QFEEVTVQQAPLTEHDGS----PDKTLEVAVRLLPRQVDHVAPVLEELIHEIVEQSQKGNWRHLVFFFFLD
Nematolebias whitei           (1) ----------MSMRGAVVGCVVLACLAVLLVERTEG HITFFSPKEILLMK--E---REGKK--DTEPRSEDG-QFEEVTIQQAPRMERDGN----PENPLKLDIRRSTKQLDHMDPLLNQIIHDIVEQHQKAR-----------
Oryzias latipes               (1) ----------MSIRGAVGVCVVLVSLMAFLVERSQG HITFFSPKELLHMRLQE---QQEKK--DMKPRSEE---FDEVILEQVPQRDAN------TDKTLELGVRFSPRQLTYVAPVIKEIIHEIVEEHQKAK-----------
Oryzias melastigma            (1) ----------MSIRGAVGVCVVLVCLMAFLVERSQG HITFFSPKELLHMRLQD---QQGKK--DMKPRSEE---FDEVILEQVPQMEAN------NDNTLELGVRLSPRQLNYVAPVIKEIIHEIVEEHQKAK-----------
Melanotaenia boesemani        (1) ----------MSMRGAVAGCVVLTCLLALLAERTDG HITFFSPKEMMLMK--E---REGKK--DIEPRSEDG-QFEEVAIQQVPQMEHNGN----PDKTLEIAVRLSPKQRNHVAPMLEEFIHEIVEDNQKAK-----------
Hippoglossus hippoglossus     (1) ----------MSMRGAVAGCLVLVCLVALLAERTEG HITFFSLKEMKQMRDR-----EGTK--DLGPRSEDS-QLEPVTVPELPQVEHDGNPVSAHDKTVEFTVRLSPKQLDHVAPVLEEYIEEILEEKAQ-----------
Hippoglossus stenolepis       (1) ----------MSMRGAVAGCLVLVCLVALLAERTEG HITFFSLKEMKQMRDR-----EGTK--DLGPRSEDS-QLEPVTVPELPQVEHDGN--P--DKTVEFTVRLSPKQLDHVAPVLEEYIEEILEEKAQ-----------
Pleuronectes platessa         (1) ----------MSMRGAVAGCLVLVCLVALLAERAEG HITFFSLKEMKQMRDR-----EGTK--DLGPRSEDS-QLEPVTVPELPQVEQDGNPVSSQDKTVEFTVRLSPKQLDHVAPVLEEYIEEILEEKAK-----------
Takifugu rubripes             (1) ----------MSMRGAVTGCLVLVCLLVLLAERAEG HITFFSPKEMMVLKQE----QEGKK--FVEPRSEDE-QFEEVTVQQLPQDKT--------DKTVAIAIHLSPQQLDHAVPVLEEIIKEMVEKAK-----------
Notolabrus celidotus          (1) ----------MSMRGAVAGCVVLVCLVALLAERAEG HITFFSPKEMMLMK--E---REGRK--DMEPRSEDL-QFEEVTAQQPPQ--E--------KTVEIAIRLSPKQLDQVAPVLEEIIHEIAEEHQKAK-----------
Cheilinus undulatus           (1) ----------MSMRRAVAGCVVLACLVALLAERTEG HITFFSPKEMMLMK--E---REGRK--DMEPRSEDG-HFEVVTVQLPSQVEHVGN----TDKGVDISVHIPP---NQVAPMLEEIMDEIGEEHQRAK-----------
Sebastes umbrosus             (1) ----------MSMRGAVAGCLVLACLLALLAERTDG HITFFSPKEMMLMK--E---REGRK--DMEPRSEDG-QFEEVTVQLP-QVEHVGN----PDKTVEIAVRLSPQQLDHVAPLLEDIIHEIVEDHQQAK-----------
Epinephelus fuscoguttatus     (1) ----------MSMRGAVAGCLVLACLVALLAERTDG HITFFSPKELMLMK--E---REGRK--DMEPRSEDG-QFEEVTVQQLPLVEGGGN----HDKTVEIAVHLSPQQLDHVAPVLEEIIHGIVEERQKAK-----------
Epinephelus lanceolatus       (1) --------- -MSMRGAVAGCLVLACLVALLAERTDG HITFFSPKELMLMK--E---REGRK--DMEPRSEDG-QFEEVTVQQLPLVEGGGN----HDKTVEIAVRLSPQQLDHVAPVLEEIIHEIVEERQKAK-----------
Epinephelus moara             (1) ----------MSMRRAVAGCLVLACLVALLAERTDG HITFFSPKELMLMK--E---REGRK--DMEPRSEDG-QFEEVTVQQLPLVEGGGN----HDKTVEIAVRLSPQQLDHVAPVLEEIIHEIVEERQKAK-----------
Etheostoma cragini            (1) ------------MRGAVAGCLVLACLVALLVEKTEG HITFFSPKEMMLFK--E---REGRK--DMEPKS-NG-QYEEETGQQLPRVERGES----PDETLEIGVRLSPKQLD----VLKENIHEIVEEHQKAK-----------
Perca fluviatilis             (1) ----------MSMRGAVAGCLVLACLVALLAERTEG HITFFSPKEMMLMK--E---REGRK--DMEPRS-NG-QFEEDAVQQLSRVERGEN----PDETVEIGVRLSPKQLDHVALVLKEIIHEIVEEHQKAK-----------
Sander lucioperca             (1) ----------MSMRGAVAGCLVLACLVMLLAERTEG HITFFSPKEMMLMK--E---REGRK--DTEPRL-NG-QFEEDRVQQLLRVEHGEN----PDKTVEIGVRLSPKQLDRVAPVLKEIIHEIVEEHQKAK-----------
Gymnodraco acuticeps          (1) ----------MSMRGAVTGCLVLACLVVLLAERTEG HITFFSPKEMMLMK--E---KEGRK--DMEPRSEDG-QFEEVTVQQLPEVEHG-N----PDKTVEIAIRLSPEQLVQVAPVIEEIIHDIVEEHQKAK-----------
Pseudochaenichthys georgianus (1) ----------MSMRGAVTGCLVLACLVVLLAERTEG HITFFSPKEMMLMK--E---REGRK--DMEPRSEDG-QFEEVTVQQLPEVEHG-N----PDKTVEIAIRLSPEQLAQVAPVLEEIIHDIVEERQKAK-----------
Trematomus bernacchii         (1) ----------MSMRGAVTGCLVLACLVVLLAERTEG HITFFSPKEMMLMK--E---REGRK--DMEPRSEDG-QFEEVTVQQLPEVEHD-N----PDKTVEIAIRLSPEQLAQVAPVLEEIIHDIVEEHQKAK-----------
Chelmon rostratus             (1) ----------MSMRGAVAGCMVLVCLVALLAERTEG HITFFSPKEMMLMK--E---REGRK--DMEPRSEDG-QLEEVTVQQLPQAERGGN----PDKTVEIAIRLSPKQLDHVVPVLEEIIHEAVEERQKAK-----------
Dicentrarchus labrax          (1) ----------MSMRGAVAGCVVLVCLVALLAERTEG HITFFSPKEMMLMK--E---REGRK--DMEPRSEDG-QFEEVTVQQIPQAERGGN----PDKTVEIAIRLSPKQLDHVAPVLEEIIHDIVEERQKAK-----------
Morone saxatilis              (1) ----------MSMRGAVAGCVVLACLVALLAERTEG HITFFSPKEMMLMK--E---REGRK--DMEPRSEDG-QFEEVTVQQLPQAERGGN----PDKTVEIAIRLSPKQLDHVAPVLEEIIHDIVEERQKAK-----------
Siniperca chuatsi             (1) ----------MSMRGAVAGCLVLVCLVMLLAERTEG HITFFSPKEMMLMK--E---REGRK--DMEPRSEDG-QFDEVTVQQLPQAERGGN----PDKTVEIAVRLSPKQLDHVAPVLKEIIHEIVEERQKAK-----------
Lates calcarifer              (1) ---       MSMRGAVAGCVVLVCLVALLAERTEG HITFFSPKEMMLMK--E---REGRK--DMEPRSEDG-QFEEVTVQQLPQVQHGGN----PDKTVEIAVRLSPKQLDHVAPMLEEIIHDIVEERQKAK-----------
Cyclopterus lumpus            (1) ----------MSMCRAVAGCLVLACMAALLAERTEG HITFFSPKEMMLMK--E---R---K--DMEPRSEDG-QLEEVTVQQLPRLDRGGN----PDQTVEIAVRLSPKQLDHVTPVLEEILHEIEQELEKAK-----------
Gasterosteus aculeatus        (1) ----------MSMRRAVVGCLVLACLVGLLAERTEG HITFFSPKEMMLMK--E---REGRK--DMEPRSEDG-QFEEVTVQQLPRLEHGGN----PDKIVEVAVRLSPKQLDRVSPVLEDIIHDIEEELQKGENVETAS------
Pungitius pungitius           (1) ----------MSMRRAVVGCLVLACLVGLLAERAEG HITFFSPKELMLMK--E---REGRK--NMEPRSEDG-QFEEVTVQQLPLLEHGGN----PDKIVEVAVRLSPKQLDHGSPVLEEIIHDIEEELQKAK-----------
Scomber japonicus             (1) ----------MSMRRAVAGCVVLACLVALLAERTEG HITFLSPKEMMLLKQ-E---QEDRK--VMEPRSEDG-QLKEVTVQQFPQLERGGN----PDKTVEIAVRLSPKQLDHVAPVLEEIIHEIVE--------------
Thunnus maccoyii              (1) ----------MSMRRAVAGCVVLACLVALLAERTEG HITFFSPKEMMLMK--E---REGRK--DMEPRSEDG-QFEEVTVQQLPQVERGGN----PDKTVEIAIRLSPKQLDHVAPVLEEIIHEIVEERQKAK-----------
Toxotes jaculatrix            (1) ----------MSMRRAVAGCVVLACLVALLAERTEG HITFFSPKEMKLMK--E---REGRK--DMEPRSEDG-QFEEVTVQQLPQLEHGGN----PDKTVEIAVRLSPKQLDHVAPVLEEIIHEIVVERQKAK-----------
Micropterus dolomieu          (1) ----------MSMRGAVAGCLVLACLVAQLAERTEG HITFFSPKEMMLMK--E---REGRK--DVEPRSEDG-QFEEVTVQQLLQVERGGN----PGKTVEIALRLSPKQLDHVAPVIEEIIHEIVDERQKGVNL---------
Micropterus salmoides         (1) ----------MSMRRAVAGCLVLVCLVAQLAERTEG HITFFSPKEMMLMK--E---REGRK--DVEPRSEDG-QFEEVTVQQLLQVERGGN----PGKTVEIALRLSPKQLDHVAPVIEEIIHEIVDERQKGVNL---------
Amphiprion ocellaris          (1) ----------MSMRGAVAGCVLLVCLVALLVERTEG HITFFSPKEMKLMK--E---REGRK--DMAPRSEDG-QFEEVTVQQVPQMERSGN----QDKTVEIAVRLSPTQLDHVAPVLEEIIHEIVEHQKAK-----------
Stegastes partitus            (1) ----------MSMRGAVAGCVVLACLVALLVERTEG HITFFSPKEMKLMK--E---REGRK--DMEPRSEDG-QFEEVTVQQVPQMEHGGN----QDKTVEIAVRLTPKQLDHVAPVLEEIIHDIVDQRQKAK-----------
Scophthalmus maximus          (1) ----------MSMRGAVAGCVLLACLAALLTERTEG HITFFSPKEMMLMK--E---REGRK--DMEPRSEDG-QFEEVTVQQLPPVEHG-------DDKTVEIAVRLSPKQLDHVAPVLEEIIHEIAEKAK-----------
Haplochromis burtoni          (1) ------------MRGAVAGCVVLVCLMALLVQRTEG HITFFSPKELMLMK--E---REGRK--DMEPRSEDG-QFEEVTVQQVPQTERGGN----PEKTVEISVRLSPKQLDHVTPVLEEIIHEITQERQKAK-----------
Simochromis diagramma         (1) ------------MRGAVAGCVVLVCLMALLVQRTEG HITFFSPKELMLMK--E---REGRK--DMEPRSEDG-QFEEVTVQQVPQTERGGN----PEKTVEISVRLSPKQLDHVTPVLEEIIHEITQERQKAK-----------
Neolamprologus brichardi      (1) ------------MRGAVAGCVALVCLMALLVQRTEG HITFFSPKELMLMK--E---REGRK--DMEPRSEDG-QFEEVNVQQVPQTERGGN----PEKTVEISVRLSPKHLDHVTPVLEEIIHEITQERQKAK-----------
Oreochromis aureus            (1) ----------MSMRGAVAGCVVLVCLMALLVQRTEG HITFFSPKELMLMK--E---REGRK--DMEPRSEDG-QFEEVTVQQVPQTERGGN----PEKIVENSVRLSPKQLDHVTPVLEEIIHEITQECQKAK-----------
Mugil cephalus                (1) ----------MSMRGAVAACVVLACVMALLVERTEG HITFFSPKEMMLMK--E---REGRK--DMEPRSEDG-QFEEVTVQQVPQVERGGN----PDKTVEFAVRLSPKQLDQVAPVLEEIIHEIVEERQKAK-----------
Hippocampus zosterae          (1) ----------MSMRGAVAVCVVLACLVALLVERTQG HITFFSPKEMMLMK--E---REGRK--DMEPRSEDG-HLEDISVQQLPQAEHGGN----PEKTVDITVRLSPKQLDHVAPVLEELIHEIVDEQQKAK-----------
Myxocyprinus asiaticus        (1) ----------MSMRGAVAVCVVLACLVALLVERTQG HITFFSPKEMMLMK--E---REGRK--DMEPRSEDG-HLEDISVQQLPQAEHGGN----PEKTVDITVRLSPKQLDHVAPVLEELIHEIVDEQQKAK-----------
Syngnathus scovelli           (1) ------------MRGAVAICVVLACLVALLVERTQG HITFFSPKEMMLMK--E---REGRK--DMEPRSEDG-QFEEVAVQQLPQVEHGGN----PDKTVDITVRLSPKQLDHVAPVLEELIHEM------------------
Scatophagus argus             (1) ----------MSMRGAVAGCVLLACLVALLAESSEG HITFFSPKELRLLKQ-E---QEGRK--DMEPRSDDG-QFEEVTVQQLPQD-----------KTVEISIRLTPKQLDHVTPVLEEIIHEIVEERQKAK-----------
```

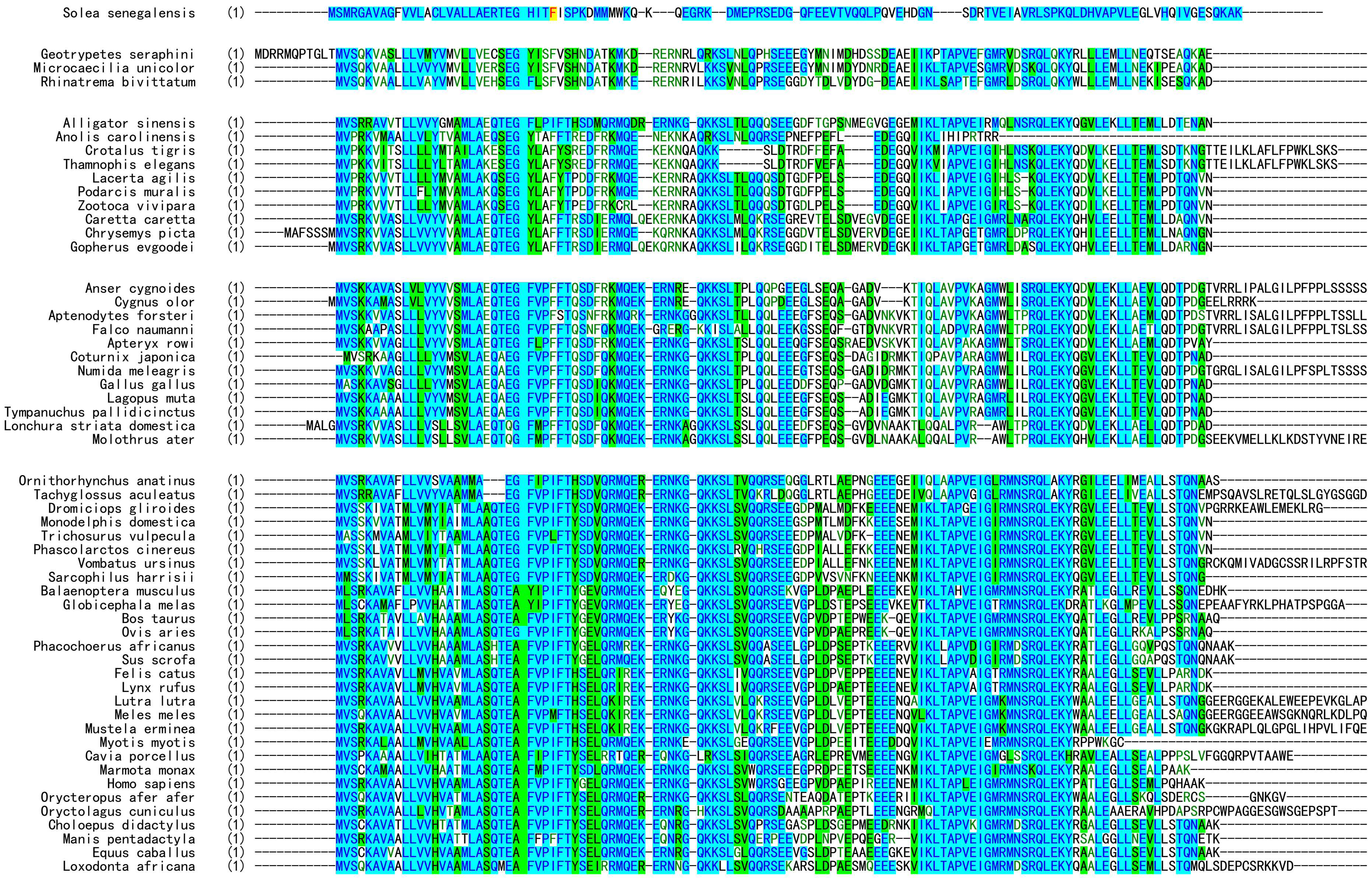

**Fig. S1.** Amino acid sequence alignment of motilin precursors from fishes to mammals. These sequences were manually downloaded from the NCBI database ([https://www.ncbi.nlm.nih.gov/gene](https://www.ncbi.nlm.nih.gov/gene)) and aligned using the software Vector NTI11.5.

```
                                   motilin precursors from H. sapiens
   NM_002418    (1) MVSRKAVAALLVVHVAAMLASQTEA  FVPIFTYGELQRMQEKERNKGQ  KKSLSVWQRSGEEGPVDPAEPIREEENEMIKLTAPLEIGMRMNSRQLEKYPATLEGLLSEMLPQHAAK-
NM_001040109    (1) MVSRKAVAALLVVHVAAMLASQTEA  FVPIFTYGELQRMQEKERNKGQ  KKSLSVWQRSGEEGPVDPAEPIREEENEMIKLTAPLEIGMRMNSRQLEKYPATLEGLLSEMLPQHAK--
NM_001184698    (1) MVSRKAVAALLVVHVAAMLASQTEA  FVPIFTYGELQRMQEKERNKGQ  KKSLSVWQRSGEEGPVDPAEPIREEENEMIKLTAPLEIGMRMNSRQLEKYPATLEGLLTAK--------

                                   motilin precursors from D. rerio
NM_001386353    (1) -MRGSVTGCVLLICVVALLAEQAEG  HIAFFSPKEMRELREKEG----  RK--DA-DSRAEGLLVDETSPEEDGGE---SAGQPVEIGLKLTAK-KSHIGSAFGKMLQNIVEEPDNAN
NM_001386354    (1) -MRGSVTGCVLLICVVALLAEQAEG  HIAFFSPKEMRELREKEG----  RK--DA-DSRAEGLLVDETSPEEDGGE---SAGQPVEIGLKLTAK-KSHIGSAFGKMLQNIVEEPDNAN

                                   motilin precursors from L. chalumnae
XM_005995467    (1) MDSRRVIGALLVVCAVVMLAERTEG  FISFFSPSDMRRMMEKEKSKAL  KKSVSLQQRSEDNSFLESPSVQYRGEGMLLKPGDTLESRVRLNMKQVDNYRDILGEMLGEILQEEQNAQ
XM_014488135    (1) MDSRRVIGALLVVCAVVMLAERTEG  FISFFSPSDMRRMMEKEKSKAL  KKSVSLQQRSEDNSFLESPSVQYRGEGMLLKPGDTLESRVRLNMKQVDNYRDILGEMLGEILQEEQNAQ
                         Signal peptide             Mature peptide                       Propeptide
```

**Fig. S2.** Amino acid sequence of motilin precursors encoded by transcript variants from *H. sapiens*, *D. rerio*, and *L. chalumnae*.
